# Disease-specific variant pathogenicity prediction using multimodal biomedical language models

**DOI:** 10.1101/2025.09.09.675184

**Authors:** Yilin Liu, David N. Cooper, Haiyuan Yu

## Abstract

Missense variants play a key role in the diagnosis of genetic disorders and in disease risk prediction. Existing methods focus primarily on the prediction of variant effects in terms of their deleteriousness, without taking into account the disease-specific context, and are therefore limited in terms of their utility in real-world diagnosis and decision making. Here, we introduce <u>di</u>sease-specific <u>va</u>riant pathogenicity prediction (DIVA), a novel deep learning framework that directly predicts specific disease types alongside the probability of deleteriousness for missense variants. Our approach integrates information from two different modalities – protein sequence and disease-related textual annotations – encoded using two pre-trained language models and optimized within a contrastive learning paradigm designed to align variants with relevant diseases in the learned representation space. Our results demonstrate that DIVA outperforms baselines and provides accurate disease predictions with high relevance to clinically curated disease annotations for missense variants. Variant deleteriousness prediction is enhanced by incorporating AlphaMissense scores through learnable weights derived from protein function annotations, which additionally boosts DIVA*’*s ability to accurately classify deleterious variants. Our work provides new insights into variant pathogenicity prediction with awareness of disease specificity, addressing a hitherto unmet need in relation to clinical variant interpretation.

## Introduction

Recent advances in next-generation sequencing technologies have led to an explosion in the number of whole genome and whole exome sequencing (WGS/WES) projects, resulting in the identification of hundreds of millions of genomic variants in human populations. As of 2022, there were more than 446,000 variants of uncertain significance (VUS) in the ClinVar^1^ database, taking up over 4 times as many as pathogenic variants (Fig. 1a, Supplementary Fig. 1). Furthermore, many genes are pleiotropic and associated with multiple clinically distinct diseases, which further significantly complicates the clinical interpretation of variants in these genes. Thorough interpretation of a variant*’*s clinical significance involves two major steps: (1) identifying deleterious variants, and (2) predicting which specific disease that each damaging variant is likely to be associated with. Previous studies have pointed out the importance of interpreting the clinical significance of variants in disease-specific settings, because classification criteria may vary both by disease and gene^2^. Despite its clinical importance, the prediction of specific disease for each damaging missense variant remains largely unexplored.

**Fig. 1.**
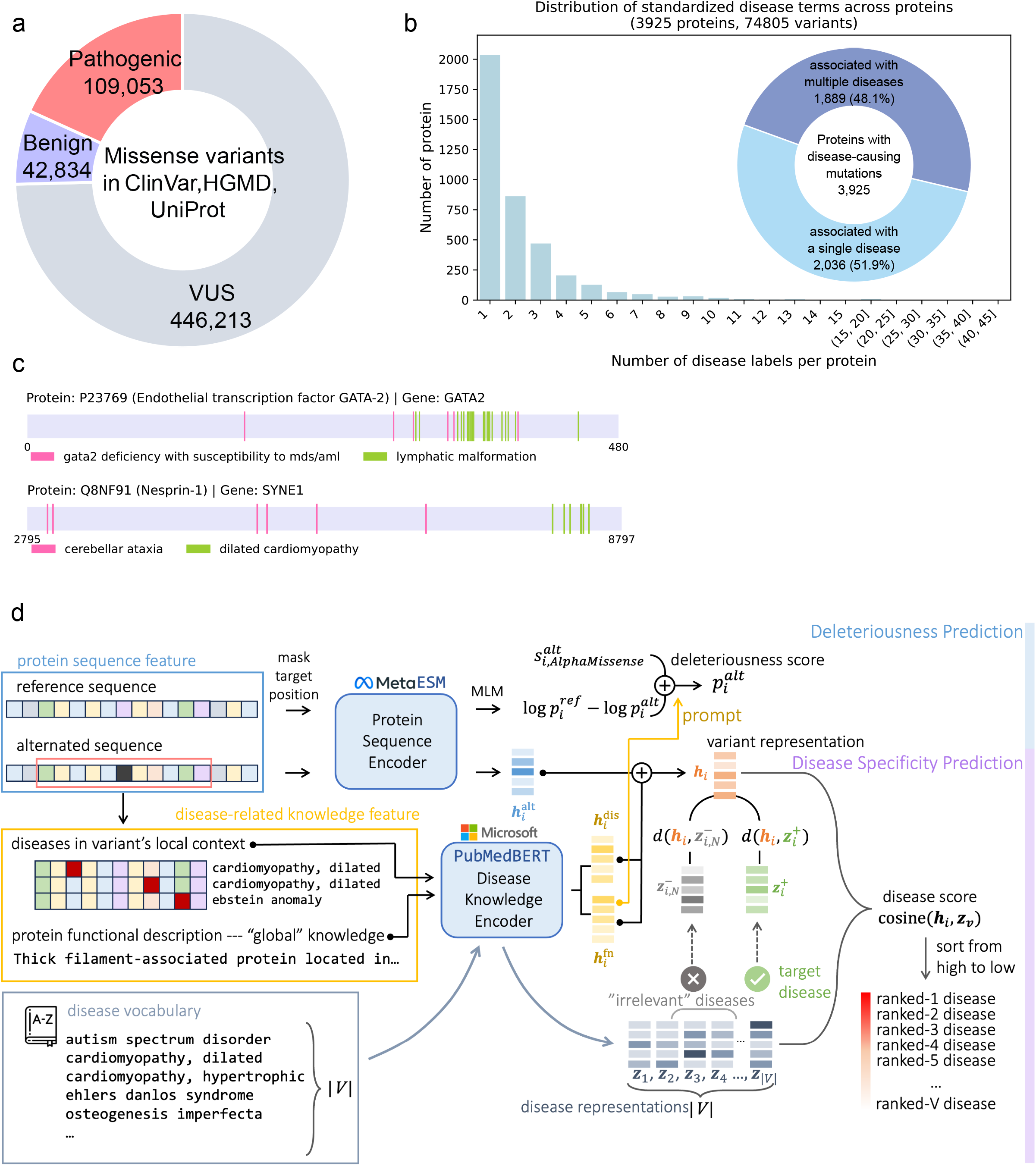
Overview of DIVA architecture. **a**, Overview of missense variants collected from ClinVar, HGMD, and UniProt databases. **b**, Summary of proteins carrying pathogenic variants with known disease conditions. The bar plot shows the distribution of the number of standardized disease terms per protein for pathogenic variants with specified diseases in the source database. The pie chart shows the number and proportion of proteins associated with single versus multiple diseases. **c**, Examples of two proteins identified with pathogenic variants leading to multiple diseases. Proteins are shown as linear representations of the protein sequences with residues of known disease mutations highlighted according to their corresponding diseases. **d**, Schematic overview of the DIVA framework. The model inputs consist of the reference and the alternated protein sequences, disease-related knowledge including the protein functional description as the global knowledge and known disease annotations of pathogenic variants located in the variant*’*s local context. It predicts the binary deleteriousness score by combining the PLM deleteriousness score with AlphaMissense score using a learned weight derived from the representation of protein functional description. For disease-specificity prediction, it additionally encodes all candidate diseases and predicts their predicted relevance to the variant according to the similarity-based disease-specificity score derived from the learned representations. The model is trained to jointly optimize the deleteriousness prediction via the binary cross-entropy loss and the disease specificity prediction using contrastive learning with hard-negative sampling (see Methods).

Numerous computational methods^3–5^ have been developed for variant effect prediction with the primary focus being placed on classifying variants as either pathogenic or benign – i.e., a binary classification of a variant*’*s deleteriousness. Recent advances in deep learning based models additionally improves variant effect prediction in terms of both accuracy and scalability. Protein language models such as the evolutionary-scale language models (ESM)^6,7^ demonstrate the capability of zero-shot inference in relation to this task, where language models pre-trained on protein sequences are directly applied to predict variant effects without any supervision from annotated variant data. AlphaMissense^8^ further integrates evolutionary data, multiple sequence alignments (MSA) and protein structure, along with the protein language model and performs large scale prediction for variants across the human proteome. Building upon these advancements, we propose a framework that leverages the ESM^9^ protein language model and AlphaMissense^8^ for their high accuracy and broad coverage in predicting the binary deleteriousness of variants. More importantly, we aim to extend beyond the binary classification of variant deleteriousness to directly predict specific diseases for each damaging variant, a clinically critical yet largely untackled task.

Despite their effectiveness in general-purpose variant effect prediction, CardioBoost^10^ and DYNA^11^ point out the issue that these methods lack specificity between genes and diseases. They improve prediction accuracy in cardiac diseases by adaptively training models using variant sequences from patients annotated with cardiomyopathies and arrhythmias. However, it is challenging to generalize these methods to genome-wide missense variants due to the lack of standardized disease annotations and challenges introduced by pleiotropic proteins (Fig. 1b,c). More importantly, none of the aforementioned tools provide any insight into what specific diseases are likely associated with a prioritized variant, which is particularly crucial for variants in pleiotropic genes.

The task of directly predicting which specific disease a deleterious variant is likely associated with is an inherently more challenging task due to the complexity and noise in submitted disease annotations for variant records in clinical databases. In addition, although variant deleteriousness can be directly inferred from protein sequences, additional information is required to establish the connection between variants and specific diseases. Recent studies^13,14^ have shown that explicitly injecting factual biological knowledge such as Gene Ontology (GO)^15^ annotations, enhances protein representation learning and improves related downstream tasks. BioBridge^16^ further demonstrates the feasibility and effectiveness of bridging multimodal biomedical knowledge in representation learning and shows its application in protein-disease association prediction. However, we argue that these methods are not sufficient for variant-level disease specificity prediction, as they tend to over-generalize across the entire protein without capturing the local context of individual variants.

In this study, we introduce <u>di</u>sease specific <u>va</u>riant pathogenicity model (DIVA), a deep learning based approach to effectively predict specific disease types along with deleteriousness for missense variants. We formulate this problem as a representation-based similarity search task, where the model generates a ranked list of most relevant diseases according to specificity scores between learned representations of the variant and all diseases. To this end, DIVA learns variant representations by leveraging contextual disease-related knowledge gathered from known mutations and protein sequence information, both of which are readily available for the majority of variants. It uses two pre-trained language models – the ESM^9^ protein language model and the PubMedBERT^17^ model – to generate initial representations of both types of information, and subsequently processes with additional lightweight modules to learn disease-specific variant representations based on the pre-trained knowledge. We further propose a contrastive learning framework with a hard negative sampling strategy to effectively train the model to distinguish between relevant and irrelevant diseases for each variant.

We evaluate DIVA for variant disease specificity prediction on a curated set of missense variants with known disease annotations. We demonstrate DIVA*’*s capability of learning a variant representation space that captures disease specificity and exhibits superior performance in variant disease specificity predictions with comparison to competitive baselines. Additionally, we benchmarked its effectiveness in variant deleteriousness prediction on our curated testing set along with the AlphaMissense cancer hotspot benchmark set. Finally, we release our extensive prediction across the large scale VUSs along with a web tool, providing a valuable resource to enhance our understanding of variant prioritization with awareness of disease specificity.

## Results

### DIVA model architecture

DIVA is a deep learning framework that extends variant pathogenicity prediction from the binary classification of deleteriousness to directly predicting specific diseases that are most likely associated with each prioritized damaging variant (Fig. 1d). It integrates state-of-the-art models – ESM^9^ and AlphaMissense^8^ – to enable accurate variant deleteriousness scoring, and further potentiates disease-specific prediction by incorporating variant contextual information from disease-related knowledge and protein sequence. The main architecture contains two pre-trained language models as core encoding components - the protein language model ESM^9^ to encode protein sequence and a biomedical domain-specific language model PubMedBERT^17^ to process disease-related knowledge. The model learns to aggregate these initial representations and compute a similarity score between each variant and candidate diseases. Based on these scores, it produces a ranked list of diseases with the top-ranked entries considered to be most likely associated with the variant. Through a contrastive learning with hard-negative sampling strategy, it is effectively optimized to bring each variant closer to its true disease labels while further away from unrelated diseases in the learned representation space.

To enable the model to directly predict specific disease for each prioritized variant, we further incorporate disease-related knowledge in addition to the protein sequence as context features. Motivated by the observation that disease-causing mutations are likely to form clusters along the protein (Fig. 1c), we collect disease annotations from variants residing in the local sequence neighborhood as local disease context features. In addition, we incorporate the functional description texts for each corresponding protein to represent the global functional context of the variant, which captures the essence that disease etiology is closely related to protein function in biological processes. By encoding disease-related knowledge through the PubMedBERT model and aggregating with ESM-generated variant sequence representations, the model learns the final variant representation. To measure the relevance of all candidate diseases to a given variant, we project representations of both the variant and all candidate diseases into a shared representation space and define a disease specificity score based on their proximity. Specifically, we use the cosine similarity between the representations of the variant and each candidate disease, with higher scores indicating stronger predicted associations between the variant and the disease.

DIVA employs a distinct contrastive learning strategy to push variants closer to their relevant diseases in the shared representation space. We use the information noise-contrastive estimation (InfoNCE) loss function^19^ which is designed to enforce separation of positive and negative samples in the representation space and inherently flexible to handle multiple negative samples. For each variant, we intentionally select hard negatives as irrelevant disease terms that receive high disease specificity scores in the model*’*s latest prediction, reflecting those that currently cause the most confusion to the model. In particular, we exclude disease terms from negative selection if they are semantically similar to the positive disease label, thereby preventing closely related terms such as disease synonyms from being mistakenly treated as negatives. In addition, we use the binary cross entropy loss for variant deleteriousness prediction, allowing the model to learn a weighting mechanism that adaptively enhances ESM-produced deleteriousness score using AlphaMissense prediction. Altogether, the contrastive loss and the binary loss are combined to jointly optimize the DIVA framework, enabling it to directly predict specific diseases associated with each variant, in addition to assessing its deleteriousness.

In practice, we realized an inherent challenge that clinically available disease-annotated variants are often insufficient in both quantity and quality to support comprehensively training both models from scratch. Therefore, in the actual training process, we focused on optimizing the projection and aggregation modules and maintained most parameters in ESM and PubMedBERT. Specifically, we froze all parameters in PubMedBERT and fine-tuned only the last hidden layer of the ESM model, thereby preserving the domain-specific knowledge encoded in PubMedBERT pre-trained weights while enabling adaptation to our task in a computationally efficient manner.

### DIVA accurately predicts disease specificity while maintaining strong performance in binary deleteriousness classification

Even though we integrated ESM and AlphaMissense for variant deleteriousness scoring, we still conducted comprehensive evaluations to thoroughly assess the model*’*s overall performance in predicting variant pathogenicity at two levels: (1) the binary classification of variant deleteriousness, and (2) direct prediction of specific diseases associated with the target variant. At the first level, we performed the evaluation on our curated test set as well as the cancer hotspot mutation set used by AlphaMissense. We excluded all variants that were seen in our training and validation sets, ending up with 11,500 benign and 11,621 pathogenic variants in our test set, and 1,693 benign and 669 pathogenic variants in the cancer hotspot set. We used AlphaMissense and ESM zero-shot predictions as benchmarks to compare against our model, using precision and recall as primary metrics (Fig. 2a,b). We showed that DIVA achieved comparable or even better performance to these state-of-the-art models in classifying variant deleteriousness.

**Fig. 2.**
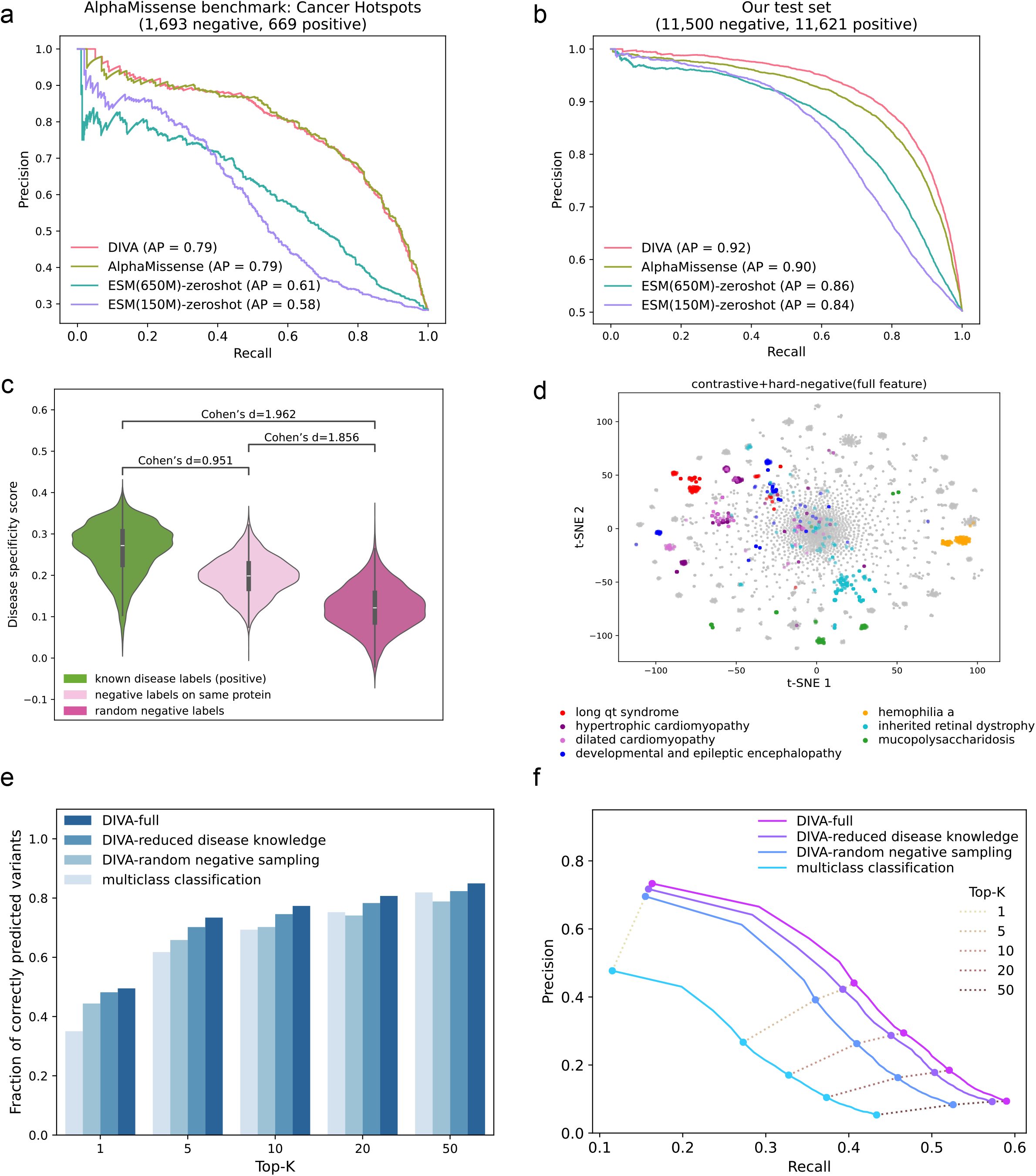
DIVA accurately predicts variant disease specificity while maintaining strong performance in deleteriousness classification. **a∼b**, Comparison of DIVA against state-of-the-art models, AlphaMissense and ESM (650M and 150M parameter settings), for binary deleteriousness prediction of variants on two data sets: **a** for AlphaMissense cancer hotspot benchmark set, and **b** for test variants curated from ClinVar. **c**, Distributions of predicted disease-specificity scores by DIVA across variants in the test set. Scores are shown for known disease labels, negative disease labels on the same protein, and random negative disease labels. **d**, t-SNE visualizations of variant representations learned by DIVA. 6 clinically distinct diseases are selected as examples and their corresponding variant representations are highlighted in different colors. **e∼f**, Assessments of DIVA-full model against ablated models for their top-K (K=1, 5, 10, 20, 50) disease predictions. **e** shows fraction of variants whose positive disease labels are correctly predicted, and **f** shows precision and recall.

We further evaluated DIVA*’*s disease specificity prediction on the 12,177 pathogenic variants in the test set with known disease labels. We first investigated DIVA for its ability to distinguish relevant diseases from unrelated ones for each variant. We examined the individual distributions of predicted disease specificity scores across three different categories of disease labels, each representing a different level of relevance to target variants. For each variant, we used pre-specified disease conditions as known disease labels, corresponding to positive labels in the classification setting. We defined irrelevant disease labels under two different settings: (1) random disease labels selected from the entire vocabulary, representing broadly unrelated diseases; and (2) protein-level unrelated disease labels, sampled within the same protein but caused by different variants. In particular, we consider the second setting more challenging, yet more reflective of real-world scenarios, where only a limited set of diseases associated with the specific gene or pathway is considered. We kept proteins with at least 5 different disease terms to allow the generation of protein-level irrelevant disease labels and to ensure fair comparisons. We investigated the distribution of model predicted disease-specificity scores for each individual category, and computed the Cohen*’*s *d*^20^ effect size metric for pairwise comparisons across all categories under the paired sample settings. As shown in Fig. 2c, DIVA consistently achieved Cohen*’*s *d* greater than the conventional threshold of large effect (*d* ≥ 0.8)^20^ across all pairwise comparisons. More impressively, it obtained Cohen*’*s *d* values of 1.962 and 1.875 when comparing known disease labels to random and protein-level unrelated disease labels respectively, both exceeding the threshold for a *“*very large*”* effect (*d* ≥ 1.2)^21^ in the expanded benchmark. This highlights DIVA*’*s strong discriminative capability in differentiating disease labels with differing relevance to target variants.

We next assessed the quality of learned variant representations by DIVA. The quality of learned representation space was evaluated according to the alignment between variants and diseases – i.e. variants sharing the same disease label should be close to each other in the representation space. We applied *t*-distributed stochastic neighbor embedding (*t*-SNE)^22^ to transform variant representations to 2D space for visualization (Fig. 2d). We chose 6 clinically distinct diseases and highlighted their corresponding variants in different colors. We observed that DIVA*’*s variant representations form well-separated clusters in the visualization. This indicates that DIVA learns high-quality variant representation space where variants are aligned according to their relevant diseases.

### Contrastive learning with hard negative sampling enhances DIVA*’*s disease-specificity prediction

To demonstrate the contribution of core components in the DIVA framework, we performed ablation study with a series of ablated models, each incorporating one of the following modifications: (1) reduced disease knowledge where protein function information was excluded; and (2) replacement of hard negative sampling with random negative sampling in computing the contrastive loss. In addition, we built a multiclass classification model as a third ablated model that completely removed contrastive learning strategy. It was trained to directly classify the known disease labels as positives, and all remaining disease labels as negatives. Each ablated model was trained and evaluated using the same set of data as the full model.

In the context of variant-disease specificity prediction, we realized the challenge of data sparsity in evaluation, where each variant is typically associated with only a few disease labels out of a large disease vocabulary. Therefore, we focused on ranking-based metrics to evaluate each model*’*s ability to retrieve relevant disease labels among top predictions, which not only offers meaningful interpretations, but also better reflects the real-world scenarios where clinicians are mostly interested in a few diseases hypothesized to be most relevant to the target variant. We used top-K accuracy (K=1, 5, 10, 20, 50) to compare the relevance of top-ranked disease predictions of each ablated model against the full model, which was computed as the fraction of variants with positive disease labels successfully included in the model*’*s top-ranked diseases (Fig. 2e). As shown in the figure, the full model consistently achieved the best results in variant disease specificity prediction compared to all ablated models. Although the multiclass classification model obtained competitive results with top-50 predicted diseases, its performance dropped rapidly when fewer top-ranked diseases were included, highlighting the robustness conferred by the contrastive learning strategy.

To evaluate performance per variant while aligning with real-world usage, we used precision and recall to measure the accuracy and sensitivity of the model*’*s top-K disease predictions. In this ranking-based setting, precision was calculated as the fraction of correctly predicted disease labels within the top-K results, while recall was the proportion of all relevant disease labels that appear in the top-K predictions. To account for variability in disease terminology and potential noise in curated disease annotations, we extended positive disease labels by including their synonyms and parent terms retrieved from Mondo^23^. By doing so, we reduced penalization for semantically correct predictions expressed in alternative forms, thereby providing a more robust and meaningful assessment of the model performance. We showed precision and recall at varying values of K to evaluate different models*’* ability in correctly prioritizing relevant diseases for each variant. As depicted in Fig. 2f, DIVA consistently achieved best precision and recall, followed by the ablated model with reduced disease knowledge, validating the contribution of including protein function annotation as global functional knowledge. We observed that models formulated as representation-based similarity search using contrastive learning strategy outperformed the multiclass classification baseline, with a notable gap observed in both metrics over the full range of top-K values. Furthermore, we noticed that hard-negative sampling led to additional performance gains compared to random negative sampling, underscoring the benefits of incorporating more informative negative disease labels in the training process.

In all experiments, we trained and evaluated DIVA using ESM (150M) as the protein sequence encoder. We further benchmarked the model using ESM of varying parameter sizes as protein encoders. Larger models yielded modest improvements in performance but incurred substantially higher training costs. Detailed comparisons of predictive performance and training efficiency is provided in Supplementary Fig. 2.

### Disease semantic analysis highlights DIVA*’*s superior performance in disease specificity prediction

We evaluated DIVA*’*s performance in variant disease-specificity prediction against the recently published state-of-the-art model BioBridge^16^. BioBridge is a foundation model built on a multi-modal biomedical knowledge graph. It first encodes uni-modal inputs separately with pre-trained language models then projects the initial representations into the same cross-modal space for specific tasks. It is shown to be able to perform disease specificity prediction at protein level as a cross-modal retrieval task, where *“*protein*”* and *“*disease*”* are set as head and tail modalities, linked by *“*associated to*”* relation. We noticed that protein level disease specificity prediction is slightly different from the variant-disease specificity prediction in that it takes the reference protein sequence without variant information. Therefore, we injected variant information for BioBridge by replacing the reference protein sequence to the alternated one, in which we substituted the single amino acid as indicated by the target variant. Fig. 3a compares DIVA versus BioBridge in terms of fractions of correctly predicted variants using top-K (K=1, 5, 10, 20, 50) for known disease variants in the test set.

**Fig. 3.**
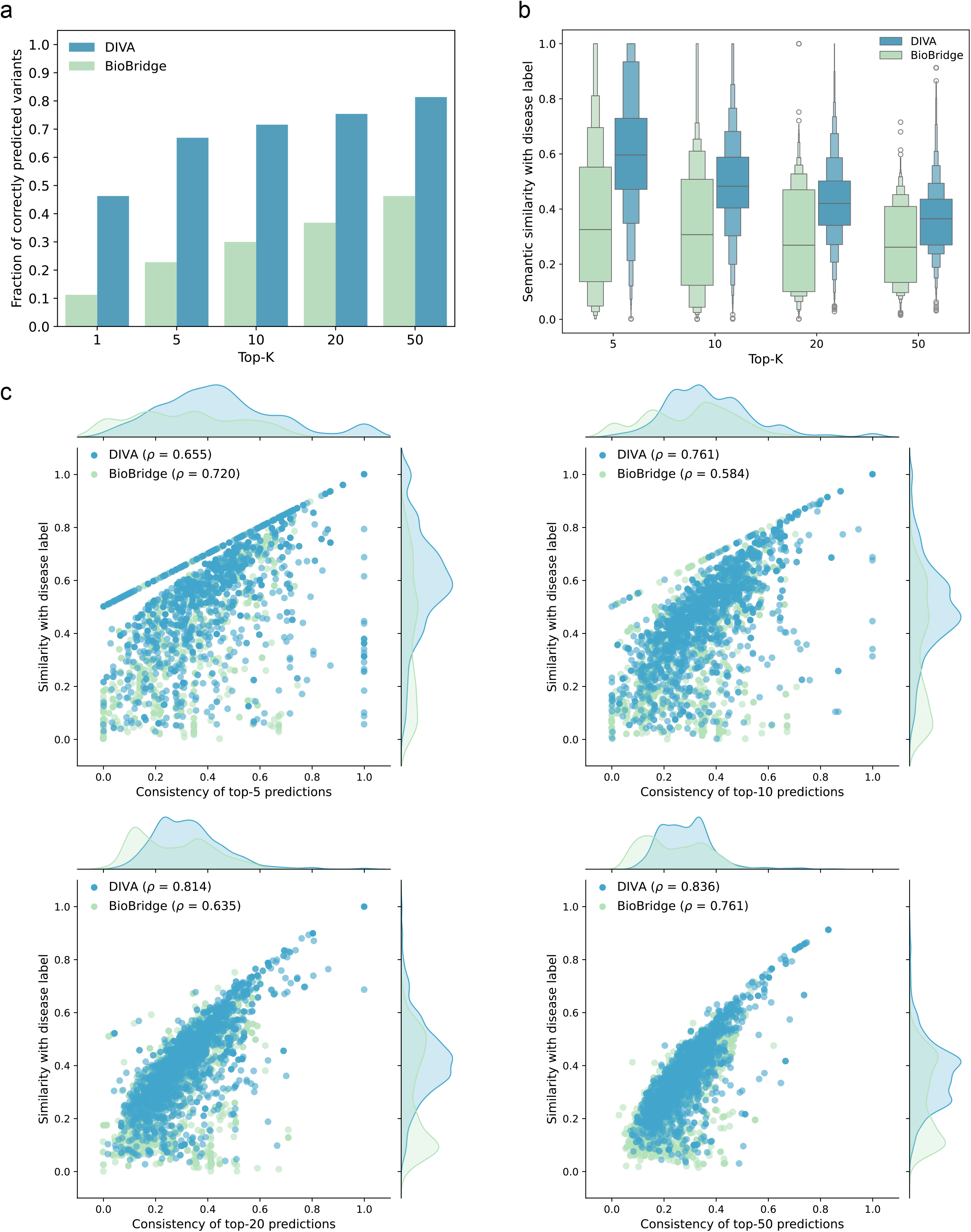
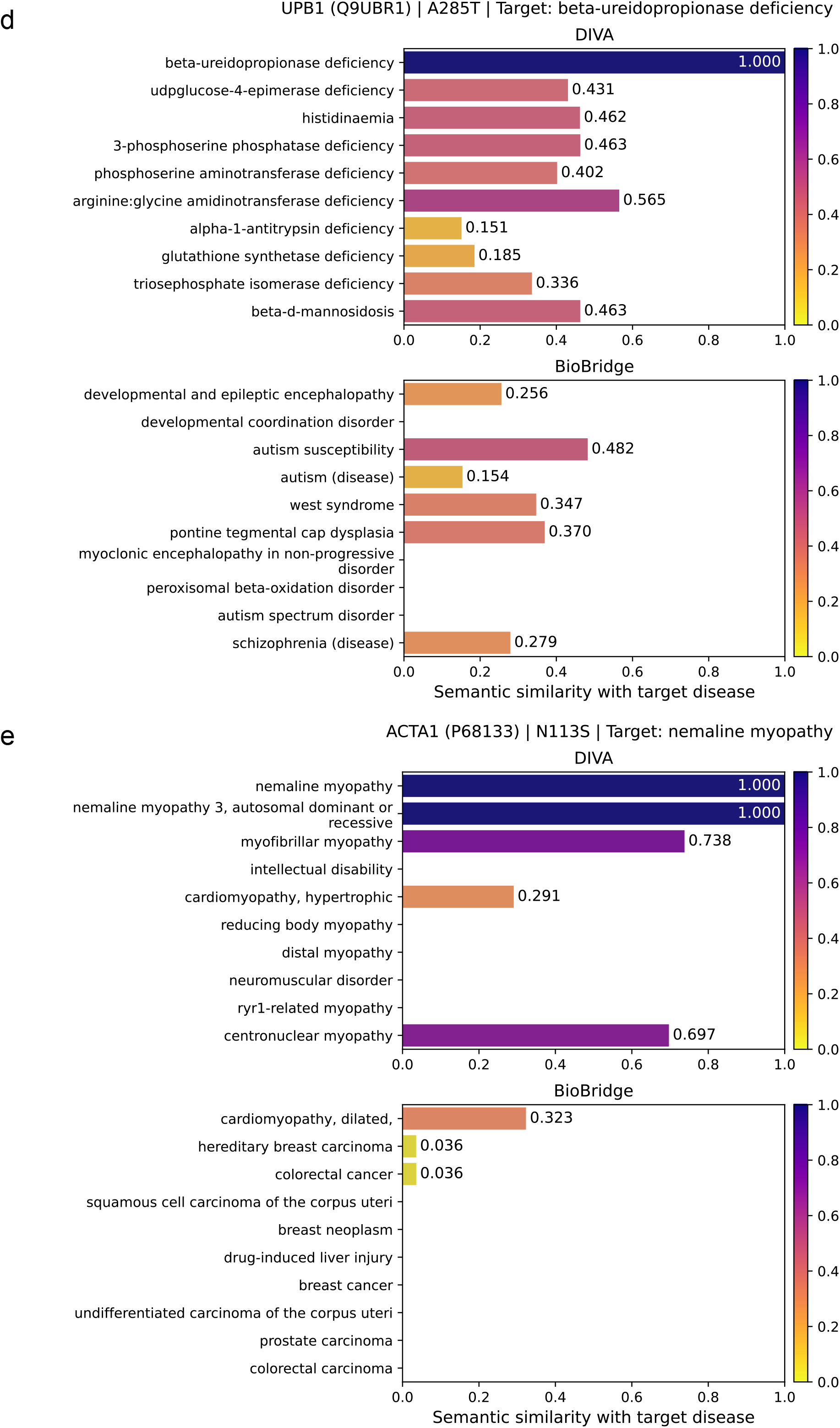
Assessment of disease-specificity prediction performance with disease semantic analysis. DIVA is compared against a state-of-the-art model, BioBridge (a multi-modal biomedical knowledge graph based foundation model) for variant disease specificity prediction. **a**, Comparison of DIVA against BioBridge in terms of the fraction of variants whose positive disease labels are correctly predicted by the model*’*s top-K disease predictions. **b**, Distribution of the average semantic similarity (aggregated per variant) between the positive disease label and each of the top-K predicted diseases. **c**, Scatter plots showing the relationships between consistency (x-axis) and correctness (similarity with disease label, y-axis) of top-K predicted diseases by DIVA and BioBridge, with marginal distributions for each metric. Pearson correlation coefficients are computed between consistency and correctness across all tested variants for each model. **d∼e**, Examples of top-10 predicted diseases for known disease variants by DIVA and BioBridge along with their semantic similarity to the target disease. Empty bars indicate semantic similarity not available for corresponding diseases. **d**, Predictions for variant A285T on UPB1 (UniProt: Q9UBR1) with disease label beta-ureidopropionase deficiency (MONDO:0013164). **e**, Predictions for variant N113S on ACTA1 (UniProt: P68133) with disease label nemaline myopathy (MONDO:0018958).

We next examined the quality of top-ranked disease predictions in terms of correctness and consistency. We evaluated correctness based on the similarity between the predicted diseases and the positive disease label, and consistency by computing similarity among top ranked disease terms predicted for the same variant. In both analyses, we used ontology-based disease semantic similarity, where we additionally retrieved corresponding phenotypes for each disease and calculated the average Lin similarity^24^ between best-matched phenotypes associated with each pair of diseases (Methods).

We first investigated the correctness of top-5, 10, 20, 50 predicted diseases by DIVA and BioBridge on known disease variants in the test set (Fig. 3b). By analyzing the distributions of average similarities between the positive disease label, we observed that DIVA consistently achieved higher similarity scores, indicating a stronger capability to prioritize diseases that are more semantically similar to the ground truth label.

We further compared the relationship between the consistency and correctness of predictions for DIVA and BioBridge (Fig. 3c). We observed that DIVA showed greater correlation between consistency and correctness throughout disease predictions ranging from top-10 to top-50. We also noticed a progressive increase in the correlation between DIVA*’*s disease prediction consistency and correctness increased as the range of top-ranked diseases expanded. In addition, the marginal distributions indicate that DIVA consistently obtained higher consistency between top-ranked predictions and higher similarity with the ground truth diseases. By contrast, BioBridge predictions showed more scattered patterns with lower correlation between consistency and correctness, and tended to be more diverse as indicated by lower consistency.

It is worth noting that DIVA was evaluated using our self-curated disease vocabulary, while BioBridge retained its original vocabulary. Evaluations were restricted to the subset of test-set variants whose positive disease labels were present in the both vocabularies. To account for differences in disease vocabulary, we additionally assessed both models using the BioBridge vocabulary (Supplementary Fig. 3,4).

### Large scale deployment of DIVA facilitates disease-specific clinical interpretation of all VUSs

Finally, we applied DIVA to all VUSs curated from ClinVar^1^. Among 410,032 VUS included for prediction, 270,183 have disease conditions submitted in their records (Fig. 4a). For variant deleteriousness prediction, we chose two score cutoffs that correspond to a false discovery rate (FDR) controlled at 0.1 and 0.05 on the test set of variants. As shown in Fig. 4b, 97,964 VUS are predicted as deleterious with a cutoff of FDR < 0.1 and 52,048 are predicted as deleterious with a more stringent cutoff of FDR < 0.05. Amon VUS predicted as deleterious using the cutoff that controlled FDR at 0.1, 63,960 variants were disease specified (Fig. 4c).

**Fig. 4.**
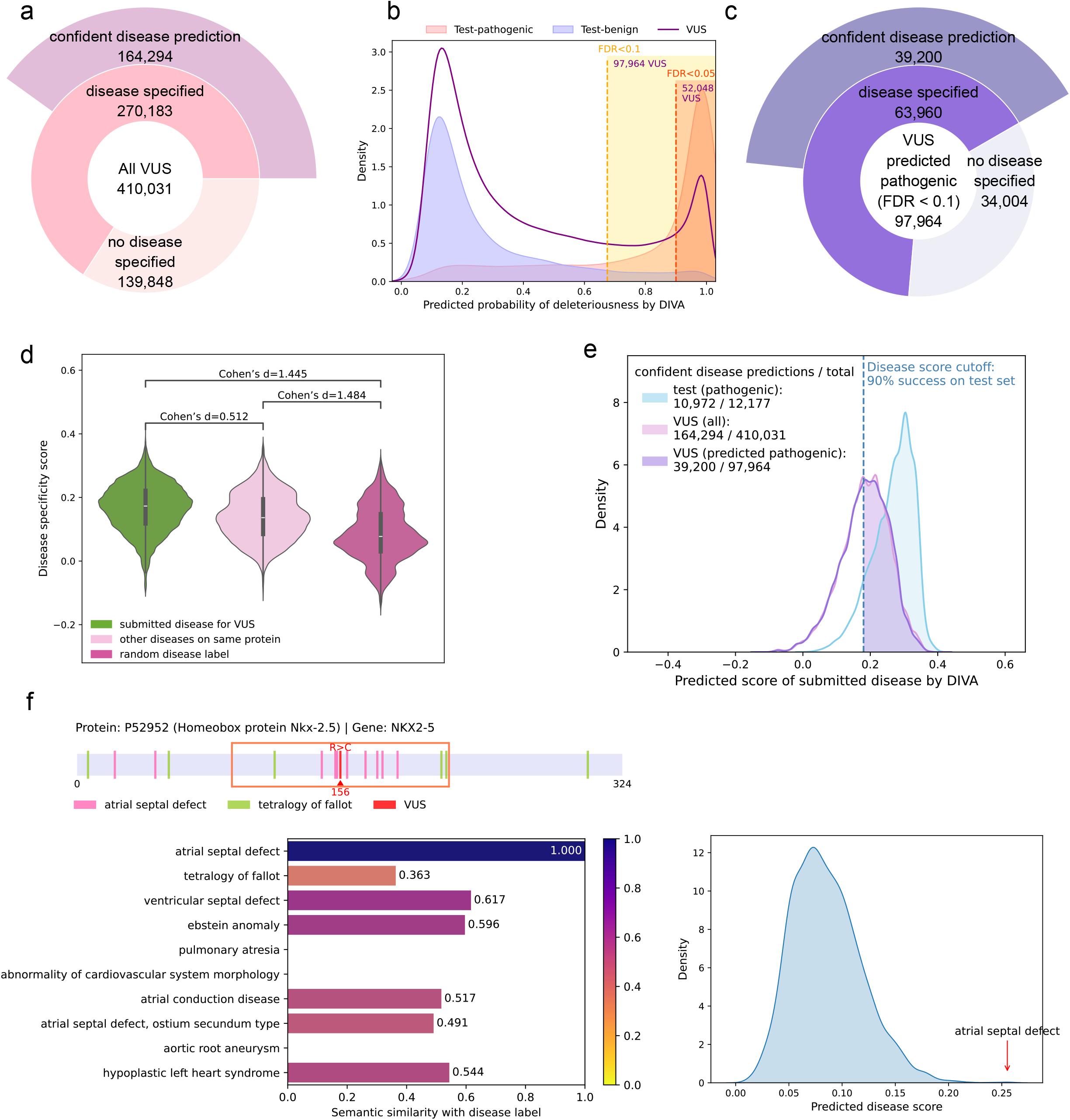
Large scale predictions by DIVA among all VUSs. We applied DIVA to 410,032 VUS collected from the ClinVar database. **a**, Summary of disease-specificity predictions on all VUS. **b**, Distributions of predicted probability of binary deleteriousness by DIVA on the test set and VUS. Two suggested thresholds are set at 0.675 to control FDR < 0.1 and 0.9 to control FDR < 0.05 at the test set, resulting in 97,964 and 52,048 VUS predicted as deleterious respectively. **c**, Summary of disease-specificity prediction on VUS predicted as pathogenic. **d**, Distributions of predicted disease-specificity score of submitted disease for VUS versus other diseases on the same protein versus random disease label chosen from the vocabulary. **e**, Distribution of predicted disease-specificity score corresponding to submitted disease conditions for variants in the test set and VUS. A disease-specificity score cutoff is set at 0.18 which yields 90% success on the test set (i.e. 90% test variants are predicted with disease-specificity score greater than 0.18 for the known disease label). **f**, An example of disease-specificity prediction for a VUS in NKX2-5 gene, for which atrial septal defect (MONDO:0006664) was submitted as the disease condition in the source database. Protein is shown as the linear representation with known disease mutations and variant context region highlighted (top). Known disease mutations are highlighted by their corresponding diseases. The red box marks the variant context region (64 residues on each side of the target position) where local disease knowledge was collected. Top-10 predicted diseases are displayed with their semantic similarity to the target disease in the bar plot (left). Distribution of predicted disease-specificity scores across all candidate diseases for the variant is shown in the density plot (right).

For disease specificity prediction, we showed the distributions of disease specificity scores for submitted disease, random disease label, and random disease on the same protein for each VUS (Fig. 4d). We curated a more comprehensive disease vocabulary for model inference by extending the training vocabulary with additional disease annotations submitted alongside VUS records. Similar to what is observed on the test set, there is still a notable difference between the distributions of submitted diseases and random disease labels, although the difference between the distributions of submitted diseases and random diseases on the same protein are much smaller for VUS. To facilitate decision making in real-world applications, we further introduced a confidence metric for variant-disease specificity prediction, analogous to the strategy applied in binary deleteriousness prediction. We set a disease specificity score cutoff to achieve 90% success in identifying true disease labels for known disease-specific variants in the test set. Applying this cutoff, 164,294 VUS were identified with confident disease predictions, including 39,200 predicted as deleterious (Fig 4. a,c,e). For each variant, we additionally define a relative confidence score, calculated as the percentile of the submitted disease*’*s predicted score against all predicted disease specificity scores across the vocabulary. Higher scores indicate a greater likelihood that the submitted disease is correctly annotated for the variant. As an example, we show DIVA*’*s disease-specificity prediction for a VUS located on the NKX2-5 gene, a pleiotropic gene found to be associated with multiple clinically distinct diseases including the atrial septal defect and tetralogy of fallot^25^ (Fig. 4f). As shown in the figure, the variant located in a local neighborhood enriched with mutations that are known to lead to the atrial septal defect. Leveraging such information, the model predicts the atrial septal defect with high confidence in its specificity score. We further examined the predicted top-10 diseases for the variant and their semantic similarity to the submitted disease. Together, these observations suggest that this VUS is likely associated with atrial septal defect.

We release all prediction results as a community resource through our web portal (https://diva.yulab.org), where users can search by protein name to retrieve variant information on the protein. On the protein details page, the web portal shows information about clinically detected variants including their distributions along the protein and DIVA*’*s prediction of disease-specificity and deleteriousness. The web portal also includes a download page with the curated disease vocabulary and all variant predictions for custom downstream analysis.

## Discussion

The recent advancement in deep learning based methods including ESM^7^ and AlphaMissense^8^, has improved variant effect prediction by bringing both higher accuracy and scalability. Despite their impressive performance, these models are designed for general-purpose prediction of variant binary deleteriousness without differentiating specific diseases or genes. While prior models such as CardioBoost^10^ and DYNA^11^ trained separate models for individual diseases to tailor variant effect prediction for specific disease contexts, they remain limited to classifying variants by deleteriousness rather than directly predicting specific disease associations. Beyond existing computational methods, we present DIVA which not only leverages disease-related context to effectively classify variant deleteriousness, but more importantly, tackles the more challenging task of directly inferring specific diseases for individual variants.

DIVA is designed to learn variant representations by leveraging contextual information from protein sequences and disease-related knowledge through two pre-trained language models. It is effectively trained under a contrastive learning framework with negative sampling strategy, while jointly optimizing the variant binary deleteriousness classification objective during the early training stage. It enables retrieval of relevant diseases through variant disease specificity scores, which are defined as the similarities between diseases and every target variant in the learned representation space. Using contextual information of disease and protein function, we additionally introduce an effective integration of AlphaMissense predictions through a learnable weighting mechanism informed by protein functional annotations. We demonstrate such strategy further enhances language model based variant deleteriousness prediction, by comparing with state-of-the-art methods on variants curated from ClinVar and cancer hotspot genes.

Through experimental analysis on missense variants with known clinically curated disease annotations, we demonstrate DIVA*’*s superior ability in producing high quality disease predictions in terms of both relevance and robustness. It shows discriminative ability in the variant-disease specificity scores for distinguishing disease labels with different levels of relevance to target variants. In addition, we validate the effectiveness of contrastive learning with hard-negative sampling strategy, which allows DIVA to obtain a high-quality variant representation space that captures variant-disease correspondences, and produces top-ranked disease predictions with high relevance to clinically curated disease labels. By employing a similarity-based scoring strategy to identify relevant diseases for each variant, DIVA inherently supports zero-shot inference on diseases unseen during training, enabling scalable application to large-scale variants across diverse disease conditions. We demonstrate this generalizability by applying DIVA to VUS for disease-specific pathogenicity prediction and releasing the results through a web-portal as a public resource to support the research community.

In conclusion, DIVA represents a new advance for variant effect prediction, offering a practical tool for clinicians in evaluating the pathogenic potential and possible disease associations for variants. By addressing the critical gap in direct inference of specific diseases at the variant level, it establishes a foundation for further research on missense variations and disease etiology.

## Methods

### DIVA model

#### Model architecture

DIVA is a multi-modal framework for disease-specific pathogenicity prediction that integrates protein sequences and disease-related text annotations (Fig. 1d). It jointly performs two tasks: the binary classification of deleteriousness and prediction of likely-associated diseases for each variant. It utilizes two pre-trained language models as core encoding components: the ESM^9^ model for protein sequences, and the PubMedBERT^17^ model for disease-related text annotations. The resulting representation vectors are passed through lightweight projection and aggregation modules that generate representations of variants informed by both sequence and disease knowledge. The model acquires disease-related knowledge from text annotations at both local and global scales. At the global scale, we retrieve descriptive texts from the *“*function*”* section of the UniProt^18^ knowledgebase of protein, denoting it as *“*global functional knowledge*”*. Local disease knowledge is derived from the local context window of 128 residues centered at the target variant along the protein sequence. Within this window, we collect known disease annotations associated with pathogenic variants in the training set. For each collected disease, we use its name, and when applicable, concatenated with its definition retrieved from Mondo^23^ as textual input. The final disease context feature is constructed by concatenating the local disease features with global functional features, separated by the *“*[SEP]*”* token as defined in the PubMedBERT model. In parallel, the ESM is applied to generate the representation of the alternate protein sequence for integration with the disease-knowledge representation, while concurrently operates the masked language modeling (MLM) objective to infer variant deleteriousness. Mathematically, the representation of variant *i* is computed as:

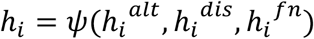

where *𝜓* denotes projection and aggregation modules applied to the initial representations of alternate protein sequence 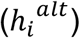, local disease feature 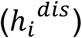, and global functional knowledge 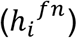.

#### Variant-disease specificity prediction

We encode all diseases in the curated vocabulary through the PubMedBERT model and project the initial representations into the same dimension as the variant representation space. Same as disease annotation features for variants, we use the concatenation of the disease name and definition retrieved from Mondo as inputs. We define variant-disease specificity score as the cosine similarity between representations of the target variant and every disease in the vocabulary. For each variant, the model ranks all diseases in the vocabulary in descending order based on their specificity scores, and thereby the top-ranked diseases are considered to be most relevant to the variant.

#### Binary deleteriousness prediction

For binary deleteriousness prediction, previous work such as ESM^18^ employs a protein language model (PLM) that operates MLM objective and infer variant deleteriousness by computing the difference of MLM logits between the reference and alternated amino acid (PLM deleteriousness score). With the release of proteome-wide variant effect prediction by AlphaMissense, we further enhance PLM-based deleteriousness prediction with AlphaMissense scores using protein-specific learnable weights. Motivated by prior studies demonstrating that protein function and disease related information facilitate variant deleteriousness prediction^9,10^, we propose to derive such learnable weights from representations of protein functional annotations which are optimized jointly within our end-to-end framework.

We enhance PLM deleteriousness score with AlphaMissense prediction by injecting a protein-specific learnable weight for each variant. As done in previous work, we pass into ESM the protein sequence with the variant position being masked and compute PLM deleteriousness score using predicted MLM logits at the masked position. Specifically, for each variant *i*, the PLM deleteriousness score is calculated as the difference between the MLM logits of the reference and alternated amino acid:

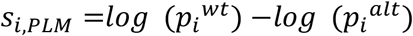

To automatically combine the PLM deleteriousness score with AlphaMissense score, we introduce learnable weights using the representation of the corresponding protein function annotation. We apply logit transformation to AlphaMissense score to align it with the scale of PLM deleteriousness score. Mathematically, for each variant *i* with alternated amino acid type *a*, the variant deleteriousness score is computed as

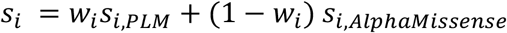

where the learnable weight 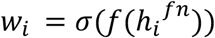 is computed by sequentially passing the representation of protein function annotation 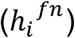 through linear projection layer *f*(⋅) and the sigmoid nonlinearity *σ* which scale the value within the range of 0 to 1.

#### Contrastive learning with hard-negative sampling for variant disease specificity prediction

We employ a contrastive learning with hard negative sampling strategy to train DIVA to effectively learn to align variants with relevant diseases in a shared representation space. We use the information noise-contrastive estimation (InfoNCE) loss function^19^ to enforce the model to learn separation of positive and negative disease labels for each variant in the representation space. In generating negative samples, we exclude diseases whose cosine similarity are ranked top-*m* (*m* = 50 in practice) with the positive disease representation, ensuring that semantically similar diseases such as disease synonyms will not be taken as negatives. Among the remaining set of diseases, we chose |𝒩| hard negatives which are disease terms that yield highest disease specificity scores for the given variants during the last iteration. Mathematically, the contrastive loss function is defined as:

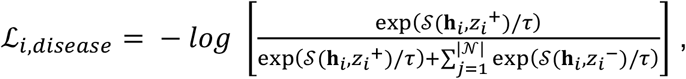

where *τ* is a temperature parameter (0.07 in our application) and 𝒮 is the similarity scoring function, which we employ as the cosine similarity.

#### Binary cross-entropy loss for enhanced deleteriousness prediction

We use binary cross-entropy (BCE) loss to fine tune the model for variant deleteriousness prediction. aiming to optimize the learnable weights we applied in combining PLM-based deleteriousness score and AlphaMissense score. The BCE loss is computed as:

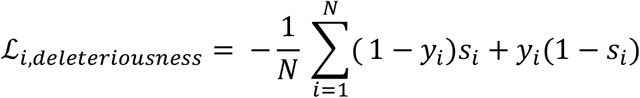

#### Model training

The model is trained with a joint loss which combines the contrastive and binary loss functions through a weighting parameter *α* that adjusts the magnitude of the deleteriousness loss.

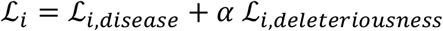

During the actual training process, we froze all parameters in PubMedBERT and fine-tuned only the last hidden layer of the ESM model. To preserve pre-trained protein knowledge encoded in ESM and mitigate overfitting, we restrict fine tuning of ESM parameters and optimization of variant deleteriousness prediction to the first two epochs. This strategy also allows the model to benefit from early adaptation to protein sequence based features while subsequently focusing on learning variant disease specificity in later training stages.

### Data preparation

#### Compilation of pathogenic, benign and VUS variants

We collected missense variants from ClinVar^1^ (version 2022/04/16), the Human Gene Mutation Database^26^ (HGMD, version 2021/02) and UniProt^18^ (2022/04 release), and categorized them into benign and pathogenic categories. For quality control purposes, we required variants from ClinVar to have review status of at least one star. Our pathogenic variants comprised ClinVar variants under the categories of *“*pathogenic*”* or *“*likely pathogenic*”*, disease-causing (DM) variants in HGMD, and variants labeled as *“*disease causing*”* in UniProt. Benign variants consisted of *“*benign*”* or *“*likely benign*”* variants from ClinVar and variants labeled as *“*polymorphism*”* from UniProt. We additionally excluded variants appeared with any conflicting clinical significance across all collected databases. After collecting missense variants from source databases, we applied Ensembl Variant Effect Predictor (VEP)^27^ to map genetic variants to UniProt identifiers and residues, resulting in 151,887 variants: 109,053 pathogenic (positive) and 42,834 benign (negative), spanning 10,652 proteins (See details in Supplementary Fig. 1). In order to train the model for deleteriousness prediction, we additionally sampled rare variants (allele frequency < 10^-3^) from the *“*controls only*”* cohort in gnomAD v2^28^ as benign variants and mapped them to variants on protein to balance the number of positive and negative samples in the curated dataset. We randomly split this dataset into 80%, 10%, 10% for training, validation, and testing sets.

#### Disease term curation

Among all assembled pathogenic variants, 97,571 are specified with disease conditions. After initial rule-based cleaning, 12,197 disease terms remained. In order to mitigate noise in disease vocabulary led by variability in disease naming and inconsistent levels of disease term granularity, we applied additional processing with reference to Mondo disease ontology^23^. We mapped each disease term to its Mondo ID when available and retrieved corresponding hierarchy from Mondo ontology. Disease terms were standardized to canonical names corresponding to Mondo IDs, and disease subtypes were merged to their parent term. The resulting disease vocabulary contains minimal redundancy while preserving most general valid terms from original records.

### Ontology-based semantic similarity

We calculated ontology-based disease semantic similarity based on disease-phenotype associations provided by the Precision Medicine Knowledge Graph (PrimeKG)^29^. PrimeKG is a data source that integrates multimodal biomedical knowledge including annotations of phenotypic abnormalities found in diseases curated by the Human Phenotype Ontology (HPO)^30^. Using the PrimeKG, we extracted phenotype annotations with HPO identifiers for each disease term. For each disease pair, we defined semantic similarity as the average Lin similarity^24^ between best-matched phenotype pairs, where each phenotype in one disease was matched to the phenotype with highest similarity in the other disease.

## Acknowledgements

The authors would like to thank Kilian Q. Weinberger, Jaehee Kim, Zizhao Zhang, and Mateo Torres for their helpful discussions. This work was supported by grants from the National Institutes of Health (NIH) RM1-GM139738, the Simons Foundation Autism Research Initiative (893926) to H.Y.

## Author Contributions

H.Y. conceived and guided the study, supervised the research and provided constructive feedback. Under close supervision of H.Y., Y.L. performed data curation, methodology development, computational experiments and analysis, and web-server construction. D.N.C. provided data sources. The manuscript was written by Y.L. and H.Y. with additional contributions and edits from D.N.C.

