## Supplementary figures for "Disease-specific variant pathogenicity prediction using multimodal biomedical language models"

Supplementary Figure 1

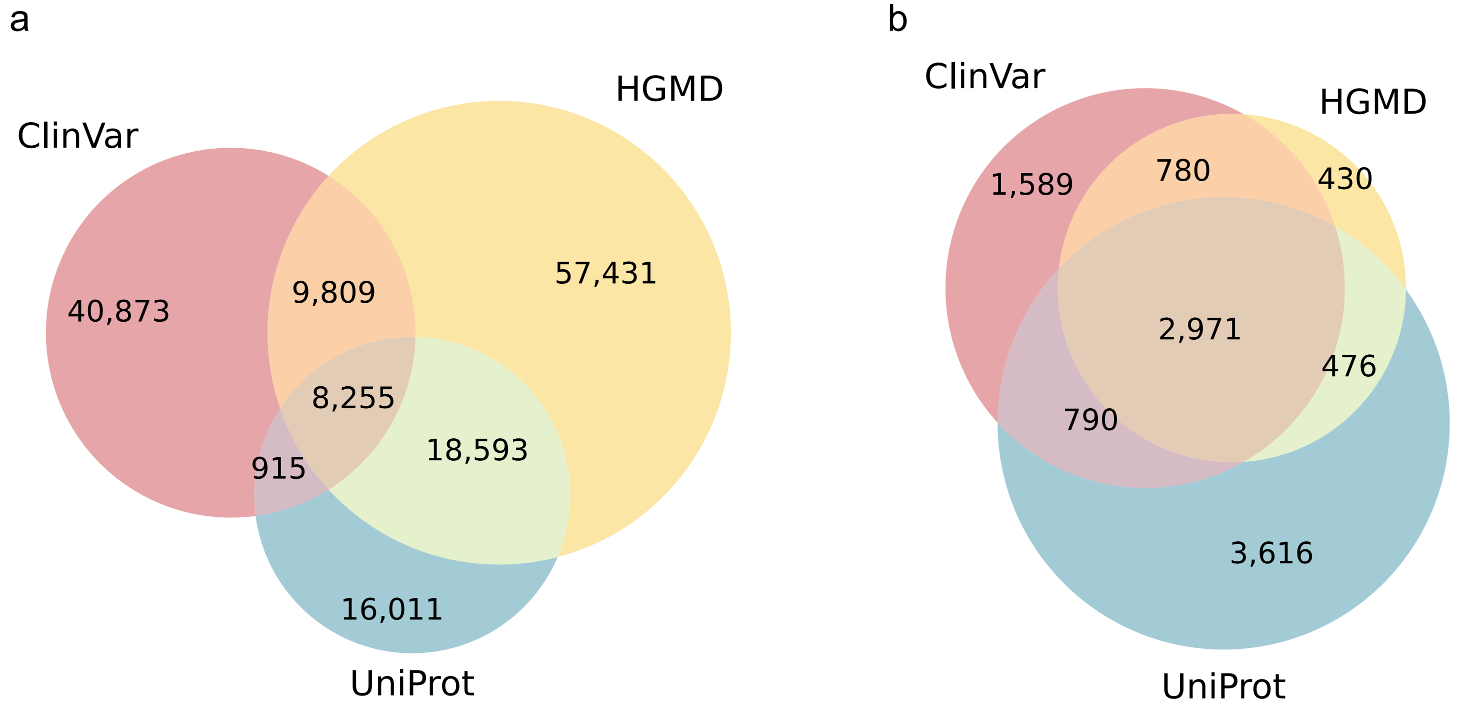


**Supplementary Figure 1 Summary of curated missense variants**
Venn diagram showing distribution and overlap of curated missense variants and corresponding proteins from the three data sources (ClinVar, HGMD, UniProt). **a**, Distribution of curated missense variants across three data sources. Numbers indicate the unique number of variants (identified by protein and residue). **b**, Distribution of proteins corresponding to curated missense variants. Numbers indicate the unique number of proteins (identified by UniProt ID).

Supplementary Figure 2

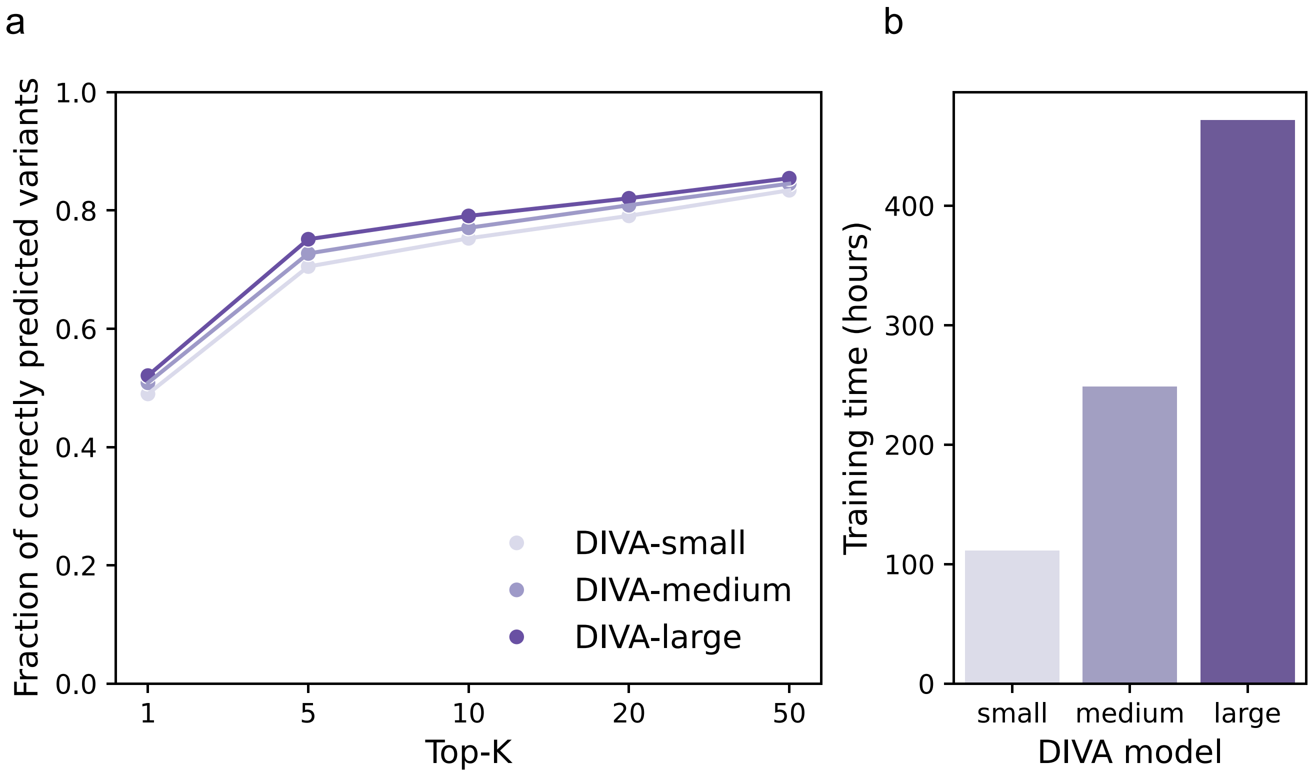


**Supplementary Figure 2 Model performance and training time under varying parameter settings.** Comparison of model performance and training time for DIVA implemented under three different parameter sizes: DIVA-small, DIVA-medium and DIVA-large. All models were trained for 20 epochs on the same dataset. **a**, Fraction of variants whose positive disease labels were correctly predicted by each model among the top-K disease predictions (K=1, 5, 10, 20, 50). **b**, Training time (hours) for each model.

Supplementary Figure 3

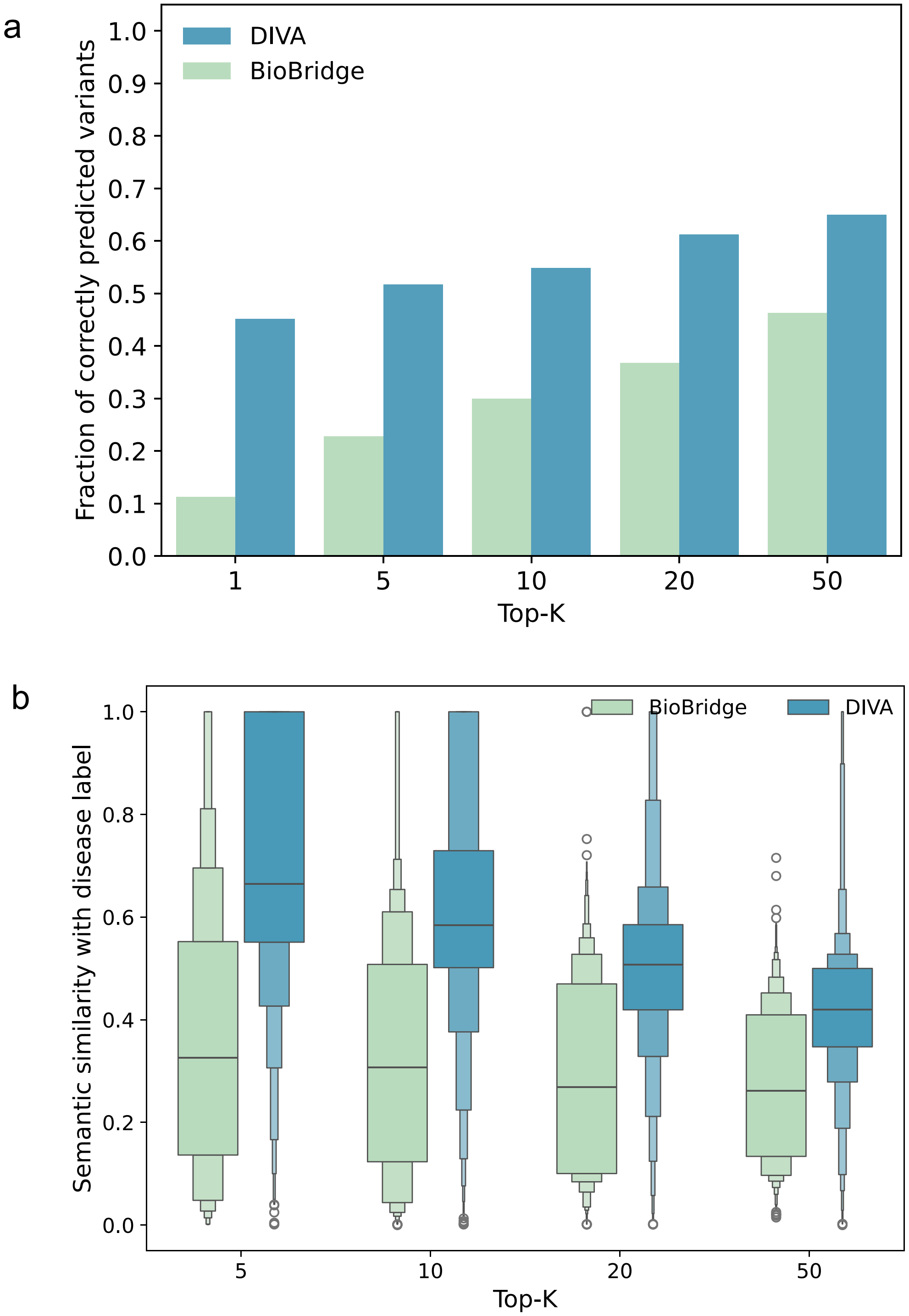


**Supplementary Figure 3 Evaluation of disease specificity prediction using BioBridge vocabulary**
Comparison of DIVA against BioBridge for disease-specificity prediction using BioBridge released disease vocabulary (17,080 terms). Evaluations were restricted to the subset of test-set variants whose positive disease labels were included in the BioBridge vocabulary. **a**, Comparison of model performance in terms of the fraction of variants for which positive disease labels were correctly predicted within the top-K disease predictions. **b**, Distribution of the average semantic similarity (aggregated per variant) between the positive disease label and each of the top-K predicted diseases.

Supplementary Figure 4

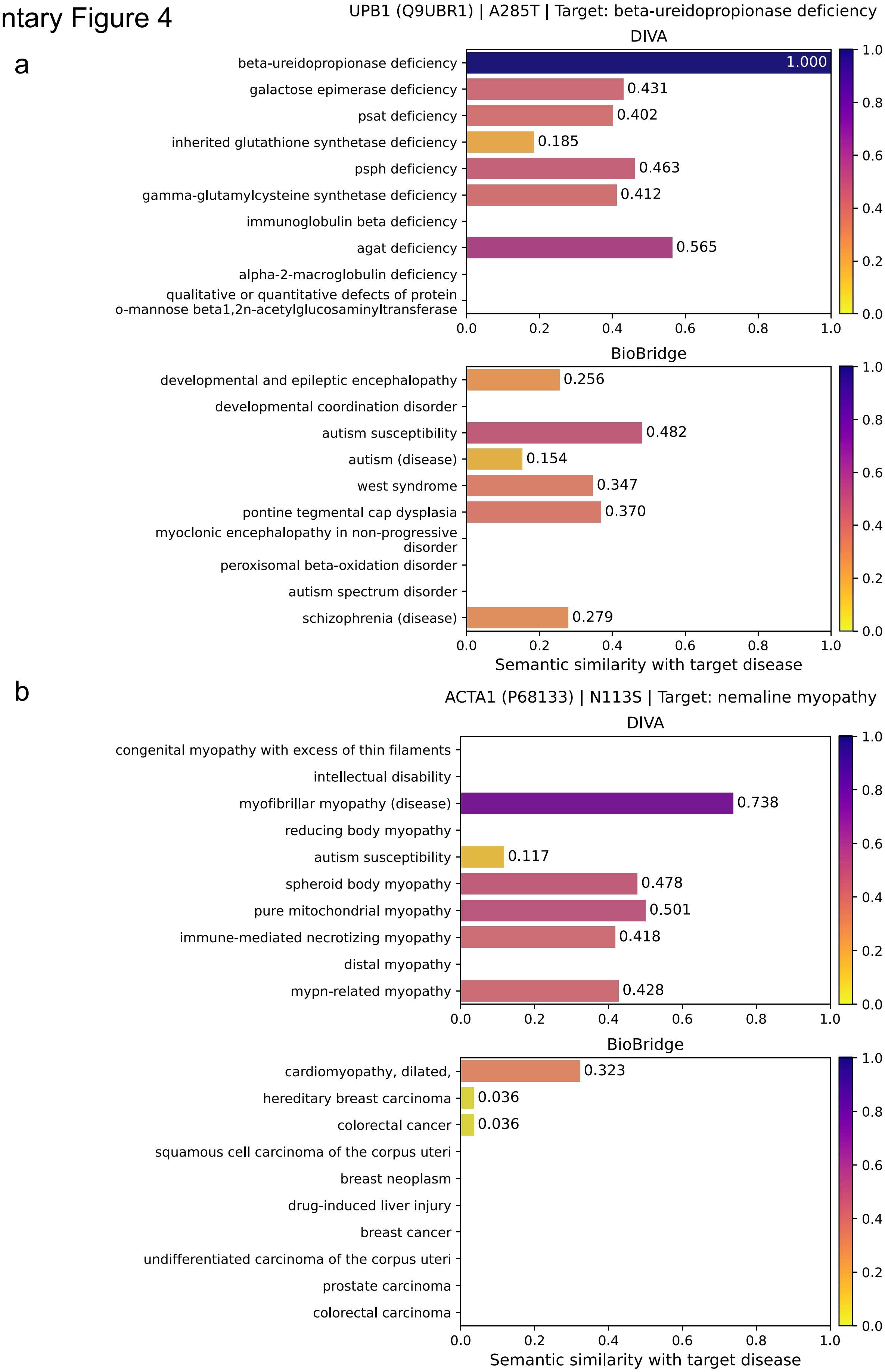


**Supplementary Figure 4 Examples of top-10 predicted diseases**

Examples of top-10 predicted diseases for known disease variants by DIVA and BioBridge along with their semantic similarity to the target disease. Predictions were made using BioBridge vocabulary for both models. **a**, Predictions for variant A285T on UPB1 (UniProt: Q9UBR1) with disease label beta-ureidopropionase deficiency (MONDO:0013164). **b**, Predictions for variant N113S on ACTA1 (UniProt: P68133) with disease label nemaline myopathy (MONDO:0018958).
